# Randomized lasso associates freshwater lake-system specific bacterial taxa with heterotrophic production through flow cytometry

**DOI:** 10.1101/392852

**Authors:** Peter Rubbens, Marian L. Schmidt, Ruben Props, Bopaiah A. Biddanda, Nico Boon, Willem Waegeman, Vincent J. Denef

**Affiliations:** KERMIT, Department of Data Analysis and Mathematical Modelling, Ghent University, Coupure Links 653, B-9000, Gent, Belgium; Department of Ecology and Evolutionary Biology, University of Michigan, 1105 North University Ave., Ann Arbor, MI 48109, USA; Department of Integrative Biology, University of Texas at Austin, 2506 Speedway, Austin, Texas 78712, USA; CMET, Center for Microbial Ecology and Technology, Ghent University, Coupure Links 653, B-9000, Gent, Belgium; Annis Water Resources Institute, Grand Valley State University, 740 West Shoreline Drive, Muskegon, MI 49441, USA

**Author notes:** Peter Rubbens and Marian L. Schmidt contributed equally to this work.

**Keywords:** bacterioplankton, 16S rRNA, flow cytometry, machine learning, variable selection, aquatic microbiology, heterotrophic productivity

## Abstract

High-(HNA) and low-nucleic acid (LNA) bacteria are two operational groups identified by flow cytometry (FCM) in aquatic systems. HNA cell density often correlates strongly with heterotrophic production, while LNA cell density does not. However, which taxa are specifically associated with these groups, and by extension, productivity has remained elusive. Here, we addressed this knowledge gap by using a machine learning-based variable selection approach that integrated FCM and 16S rRNA gene sequencing data collected from 14 freshwater lakes spanning a broad range in physicochemical conditions. There was a strong association between bacterial heterotrophic production and HNA absolute cell abundances (R^2^ = 0.65), but not with the more abundant LNA cells. This solidifies findings, mainly from marine systems, that HNA and LNA could be considered separate functional groups, the former contributing a disproportionately large share of carbon cycling. Taxa selected by the models could predict HNA and LNA absolute cell abundances at all taxonomic levels, with the highest performance at the OTU level. Selected OTUs ranged from low to high relative abundance and were mostly lake system-specific (89.5%-99.2%). A subset of selected OTUs was associated with both LNA and HNA groups (12.5%-33.3%) suggesting either phenotypic plasticity or within-OTU genetic and physiological heterogeneity. These findings may lead to the identification of systems-specific putative ecological indicators for heterotrophic productivity. Generally, our approach allows for the association of OTUs with specific functional groups in diverse ecosystems in order to improve our understanding of (microbial) biodiversity-ecosystem functioning relationships.

**Importance:** A major goal in microbial ecology is to understand how microbial community structure influences ecosystem functioning. Research is limited by the ability to readily culture most bacteria present in the environment and the difference in bacterial physiology *in situ* compared to in laboratory culture. Various methods to directly associate bacterial taxa to functional groups in the environment are being developed. In this study, we applied machine learning methods to relate taxonomic data obtained from marker gene surveys to functional groups identified by flow cytometry. This allowed us to identify the taxa that are associated with heterotrophic productivity in freshwater lakes and indicated that the key contributors were highly system-specific, regularly rare members of the community, and that some could switch between being low and high contributors. Our approach provides a promising framework to identify taxa that contribute to ecosystem functioning and can be further developed to explore microbial contributions beyond heterotrophic production.

## Introduction

A key goal in the field of microbial ecology is to understand the relationship between microbial diversity and ecosystem functioning. However, it is challenging to associate bacterial taxa to specific ecosystem processes. Marker gene surveys have shown that natural bacterial communities are extremely diverse and the presence of a taxon does not imply its activity. The taxa observed in these surveys may have low metabolic potential, be dormant, or have recently died (1, 2). An additional hurdle is that the current standard unit of measure for microbial taxonomic analysis is relative abundance. This results in a negative correlation bias (3), which makes it difficult to quantitatively associate specific microbial taxa with microbial ecosystem functions using traditional correlation measures (4). Therefore, in order to ultimately model and predict bacterial communities, new methodologies, which integrate different data types, are needed to associate bacterial taxa with ecosystem functions (5).

One such advance is the use of flow cytometry (FCM), which has been used extensively to study aquatic microbial communities (6–8). This single-cell technology partitions individual microbial cells into phenotypic groups based on their observable optical characteristics. Most commonly, cells are stained with a nucleic acid stain (*e.g.* SYBR Green I) and upon analysis assigned to either a low nucleic acid (LNA) or a high nucleic acid (HNA) group (9–12). HNA cells differ from LNA cells in both a considerable increase in fluorescence due to cellular nucleic acid content and scatter intensity due to cell morphology. The HNA group is thought to contribute more, whereas the LNA population has been considered to contribute less to productivity of a microbial community (6, 13–15). This is based on positive linear relationships between HNA abundance and (a) bacterial heterotrophic production (BP) (10, 14–17), (b) bacterial activity measured using the dye 5-cyano-2,3-ditolyl tetrazolium chloride (18, 19), (c) phytoplankton abundance (20), and (d) dissolved organic carbon concentrations (21). Additionally, growth rates are higher for HNA than LNA cells (13, 16, 22) and HNA cells accrue cell damage significantly faster than the LNA cells under temperature (23) and chemical oxidant stress (24). In contrast, LNA bacterial growth rates are positively correlated with temperature and negatively correlated with chlorophyll a (25). However, it is important to note that LNA cells are often smaller than HNA cells (12, 25–27) and therefore LNA cells could have similar amino acid incorporation rates compared to HNA cells when evaluating biomass-specific production (12).

Here we used a data-driven approach to associate the dynamics of individual taxa with those of the LNA and HNA groups in freshwater lakes by adopting a machine learning variable selection strategy. We applied two variable selection methods, the Randomized Lasso (RL) (28) and the Boruta algorithm (29) to associate individual taxa with HNA and LNA cell abundances. These methods extend on traditional machine learning algorithms (*i.e.* the Lasso and Random forest algorithm for the RL and Boruta algorithm, respectively) by making use of resampling and randomization. These extensions are needed as (a) the Lasso algorithm is not suited for compositional data because the regression coefficients have an unclear interpretation, and single variables may be selected when correlated to other variables (30), and (b) Random Forest algorithms can be biased towards correlated variables (31), which is an intrinsic issue with relative abundance data (3). The extended methods allow the user to either assign a probability of selection (RL) or statistically decide which taxa to select (Boruta).

We gathered samples from three types of lake systems (i) a set of oligo-to eutrophic small inland lakes, (ii) a short residence time mesotrophic freshwater estuary lake (Muskegon Lake), and (iii) a large oligotrophic Great Lake (Lake Michigan), all located in Michigan, USA. We then used the RL and Boruta algorithms to associate specific bacterial taxa to HNA and LNA FCM functional groups, and via the observed HNA-productivity relationship, to functioning. To validate the RL-based association with the HNA and/or LNA group, we correlated taxon abundances with specific regions within the FCM fingerprint at finer resolution (*i.e.* bins) without prior knowledge of the HNA/LNA groups. Furthermore, we tested for phylogenetic conservation of HNA and LNA functional groups using the probabilities from the RL output and for the association between the selected taxa and productivity.

## Results

### Study lakes are dominated by LNA cells

The inland lakes (6.3 × 10^6^ cells/mL) and Muskegon Lake (6.0 × 10^6^ cell/mL) had significantly higher total cell abundances than Lake Michigan (1.7 × 10^6^ cell/mL; p = 2.7 × 10^−14^). Across all lakes, the mean proportion of HNA cell counts (HNAcc) to total cell counts was much lower (30.4 ± 9%) compared to the mean proportion of LNA cell counts (LNAcc; 69.6 ± 9%). Through ordinary least squares regression, there was a strong correlation between HNAcc and LNAcc across all data (R^2^ = 0.45, P = 2 × 10^−24^; Figure 1A), however, only Lake Michigan (R^2^ = 0.59, P = 5 × 10^−11^) and Muskegon Lake (R^2^ = 0.44, P = 2 × 10^−9^) had significant correlations when the three ecosystems were considered separately.

**Figure 1:**
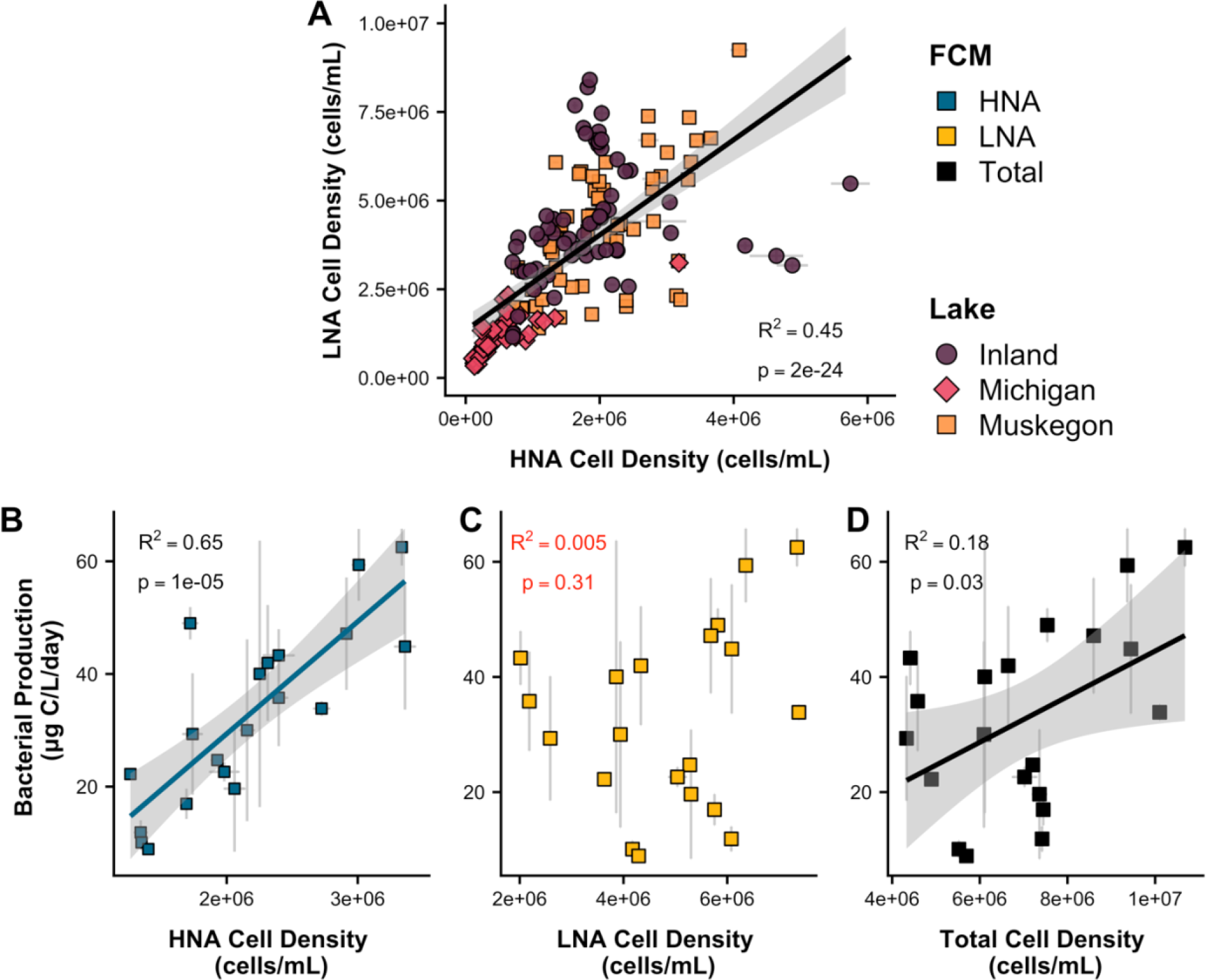
**(A)** Correlation between HNA cell counts and LNA cell counts across the three freshwater lake ecosystems. **(B-D)** Muskegon Lake bacterial heterotrophic production and its correlation with **(B)** HNA cell counts (HNAcc), **(C)** LNA cell counts, (LNAcc) and **(D)** total cell counts. The grey area in plots A, B, and D represents the 95% confidence intervals.

### HNA cell counts and heterotrophic bacterial production are strongly correlated

For mesotrophic Muskegon Lake, there was a strong correlation between total bacterial heterotrophic production and HNAcc (R^2^ = 0.65, P = 1e-05; Figure 1B), no correlation between BP and LNAcc (R^2^ = 0.005, P = 0.31; Figure 1C), and a weak correlation between heterotrophic production and total cell counts (R^2^ = 0.18, P = 0.03; Figure 1D). There was a positive (HNA) and negative (LNA) correlation between the fraction of HNA or LNA to total cells and productivity, however, the relationship was weak and not significant (R^2^ = 0.14, P = 0.057).

### Association of OTUs to HNA and LNA groups by Randomized Lasso

The relevance of specific OTUs for predicting FCM functional group abundance was assessed using the Randomized Lasso (RL), which assigns a score between 0 (unimportant) to 1 (highly important) to each taxon in function of the target variable: HNAcc or LNAcc. To assess the predictive power of a subset of OTUs based on the RL, we iteratively removed the OTUs with the lowest RL score in a recursive variable elimination scheme. 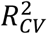, a goodnsess-of-fit measure using the *R*^2^ of how well a set of selected OTUs predicts HNAcc or LNAcc compared to true values using cross-validation, increased when lower-ranked OTUs were removed (moving from right to left on Figure 2). The increase was gradual for the inland lakes (Figure 2A) and Muskegon Lake (Figure 2C) but was abrupt for Lake Michigan (Figure 2B). The proportion of taxa that resulted in the highest 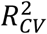 (*see solid (HNA) and dotted (LNA) lines in* Figure 2) was 10.2% of all taxa for HNA and 17.7% for LNA for the inland lakes, 4.0% for HNA and LNA for Lake Michigan, and 21.1% for both HNA and LNA in Muskegon Lake. Lake Michigan differed the most from other lake systems, having the lowest 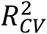, a sharp increase in 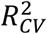 as OTUs were eliminated, and a considerably lower number of OTUs that were retained (13 for HNAcc, 10 for LNAcc). No relationship could be established between rankings of variable selection methods and the relative abundance of individual OTUs (**Figure S1**). HNAcc and LNAcc could be predicted with equivalent performance to relative HNA and LNA proportions, yet the increase between initial and optimal performance was larger (**Figure S2**). The final predictive performance was higher when relative OTU abundances were transformed using the CLR-transformation (**Figure S3**).

**Figure 2:**
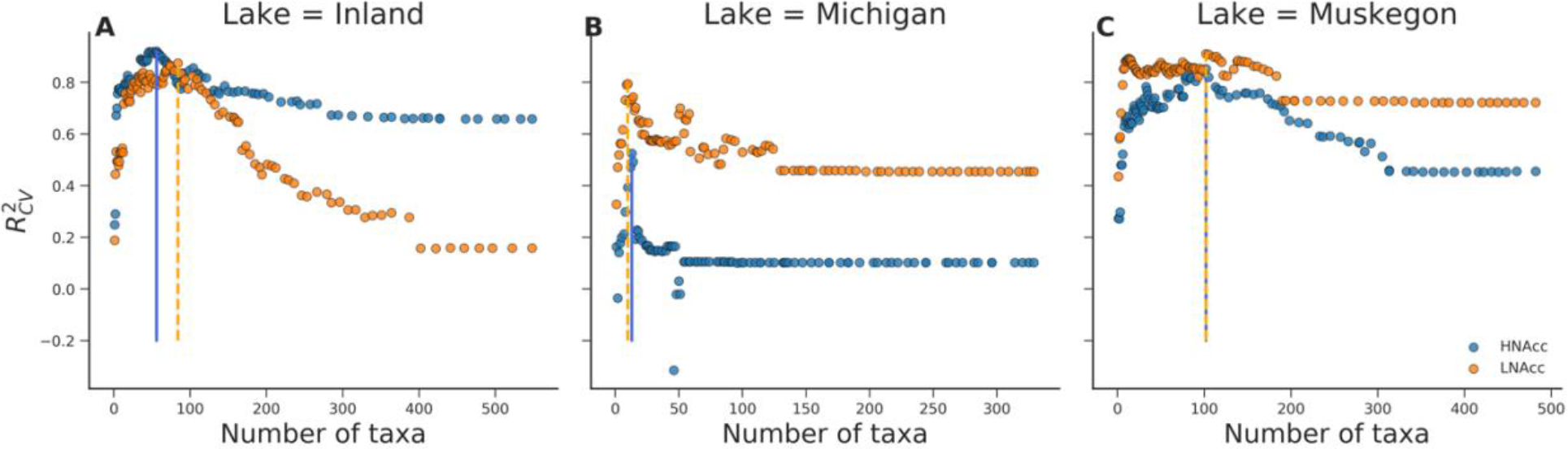
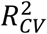 in function of the number of OTUs, which were iteratively removed based on the RL score and evaluated using the Lasso at every step. The solid (HNA) and dashed (LNA) vertical lines corresponds to the threshold (i.e., number of OTUs) which resulted in a maximal 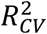. **(A)** Inland system 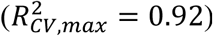, HNAcc; **(B)** Lake Michigan 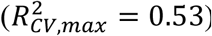, HNAcc; **(C)** Muskegon lake, HNAcc 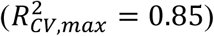; **(D)** Inland system, LNAcc 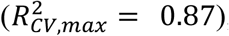; **(E)** Lake Michigan, LNAcc 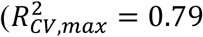; **(F)** Muskegon lake, LNAcc 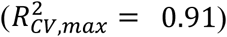.

### OTU-level predictions outperform other taxonomic levels

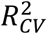 values were considerably higher than zero on all taxonomic levels, indicating that our results were consistent across all taxonomic levels and that different levels can be related to changes in HNAcc and LNAcc. While the OTU level resulted in the best prediction of HNAcc and LNAcc (Figure 3), each individual OTU had a lower RL score compared to other taxonomic levels, which on average became lower as the taxonomic level decreased (**Figure S4**). The fraction of variables (*i.e.* taxa) that could be removed to reach the maximu 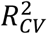 decreased as the taxonomic level became less resolved.

**Figure 3:**
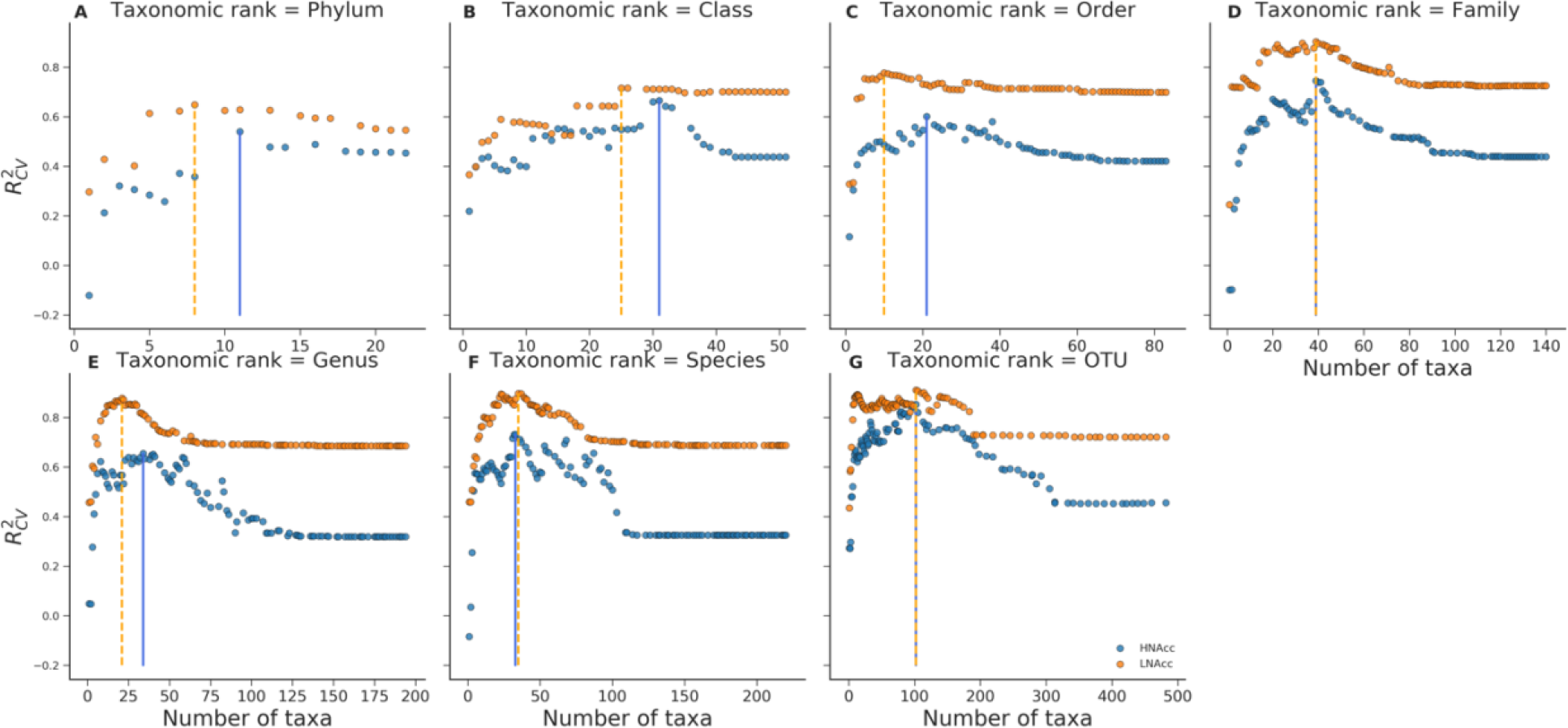
Evaluation of HNA cell counts (HNAcc) and LNA cell counts (LNAcc) predictions using the Lasso at all taxonomic levels for the Muskegon lake system, expressed in terms of 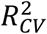, using different subsets of taxonomic variables. Subsets were determined by iteratively eliminating the lowest-ranked taxonomic variables based on the RL score.

### Validation of RL OTU selection results using the Boruta algorithm and Kendall tau statistic

Venn diagrams were constructed to visualize consistency in the number of OTUs that were selected according to the RL method, the Boruta algorithm, and individual correlations with HNAcc and LNAcc via the Kendall rank correlation coefficient (**Figure S5**). The Kendall rank correlation coefficient selected the most OTUs, followed by the RL, and then the Boruta algorithm (except for HNAcc in Lake Muskegon; **Figure S5**). The Boruta algorithm selects relevant variables based on the importance of the most permuted variable as retrieved from multiple Random Forest models (*see materials and methods*). The Boruta algorithm ranks selected OTUs as ‘1’, tentative OTUs as ‘2’, and all other OTUs have lower ranks, depending on the stage in which they were eliminated. The fraction of selected OTUs was always smaller than 1% across lake systems and functional groups (**Figure S6**). All methods agreed on only a small subset of OTUs.

For each lake system individually, the top RL-scored OTU for HNAcc was also selected by the Boruta algorithm, whereas both methods only agreed for Lake Michigan LNAcc (Table 1). Across all lake systems, OTU060 (Proteobacteria;Sphingomonadales;alfIV_unclassified) was the only OTU selected across all lake systems (LNAcc-associated). As Random Forest regressions are the base method of the Boruta algorithm, we compared the predictive power of Boruta selected OTUs to those of all OTUs using Random Forest regression. For all lake systems and functional groups, the performance increased when only Boruta-selected OTUs were included in the model (**Figure S7**). Lasso predictions, in which OTUs were selected according to the RL, were better as opposed to Random Forest predictions in which OTUs were selected according to the Boruta algorithm (**Figure S7**).

**Table 1:** Top scored OTUs according to the RL per functional population and lake ecosystem. Selection according to the Boruta algorithm is given in addition to the RL score. Descriptive statistics by means of the Kendall rank correlation coefficient have been added with level of significance in function of the HNA/LNA population. Full taxonomy of the OTUs is given in **Table S2**.

| Lake system | Functional group | OTU | RL score | Boruta selected | Kendall tau (HNA) | P-value (HNA) | Kendall tau (LNA) | P-value (LNA) | Phylum | Class | Order | Family | Genus (species) |
| --- | --- | --- | --- | --- | --- | --- | --- | --- | --- | --- | --- | --- | --- |
| Inland | HNA | OTU 369 | 0.382 | yes | -0.43 | <0.001 | -0.28 | 0.0012 | Proteobacteria | Alphaproteobacteria | Rhodospirrlalleles | alfVIII | alfVIII_unclassified |
|  | LNA | OTU 555 | 0.384 | no | 0.089 | N.S. | 0.22 | 0.011 | Proteobacteria | Deltaproteobacteria | Bdellovibrionales | Bdellovibrionacea | OM27_clade |
| Michigan | HNA | OTU 025 | 0.362 | yes | 0.46 | <0.001 | 0.41 | <0.001 | Bacteroidetes | Cytophagia | Cytophagales | bacIII | bacIII-A |
|  | LNA | OTU 168 | 0.428 | yes | 0.26 | 0.0092 | 0.4 | <0.001 | Proteobacteria | Alphaproteobacteria | Rhizobiales | alfVII | alfVII_unclassified |
| Muskegon | HNA | OTU 173 | 0.462 | yes | 0.5 | <0.001 | 0.2 | 0.019 | Bacteroidetes | Flavobacteriia | Flavobacteriales | bacII | bacII-A |
|  | LNA | OTU 029 | 0.568 | no | 0.26 | 0.0029 | 0.49 | <0.001 | Bacteroidetes | Cytophagia | Cytophagales | bacIII | bacIII-B (Algor) |

Although all methods only agreed on a minority of OTUs, we can still formulate a number of general conclusions across these methods: (1) the selected OTUs were mostly lake systems specific, (2) a small fraction of OTUs was needed to predict changes in community composition, (3) selected OTUs were associated with absolute HNA or LNA abundance, (4) top RL-ranked HNA-associated OTUs were also selected according to the Boruta algorithm and (5) when the RL and Boruta both agreed on an OTU it was always significantly correlated with both HNAcc or LNAcc.

### HNA- and LNA-associated OTUs differed across lake systems

RL-selected OTUs were mostly assigned to either the HNA or LNA groups and there was limited correspondence across lake systems between the selected OTUs (Figure 4). 1.5%-1.9% of the OTUs selected for Lake Michigan were also associated with HNAcc or LNAcc for the inland lakes or Muskegon Lake. This amount was higher for the shared OTUs between the inland lakes and Lake Muskegon, but still only amounted to 6.0% (HNAcc) or 10.5% (LNAcc) of all common OTUs. For OTUs selected in all three freshwater environments, RL scores were lake ecosystem specific, with only a significant similarity between the Inland lakes and Muskegon lake for HNAcc (r = 0.21, P = 0.0042; **Figure S8**). The Boruta algorithm selected mostly OTUs that were unique both for the lake system and FCM group (**Figure S9**).

**Figure 4:**
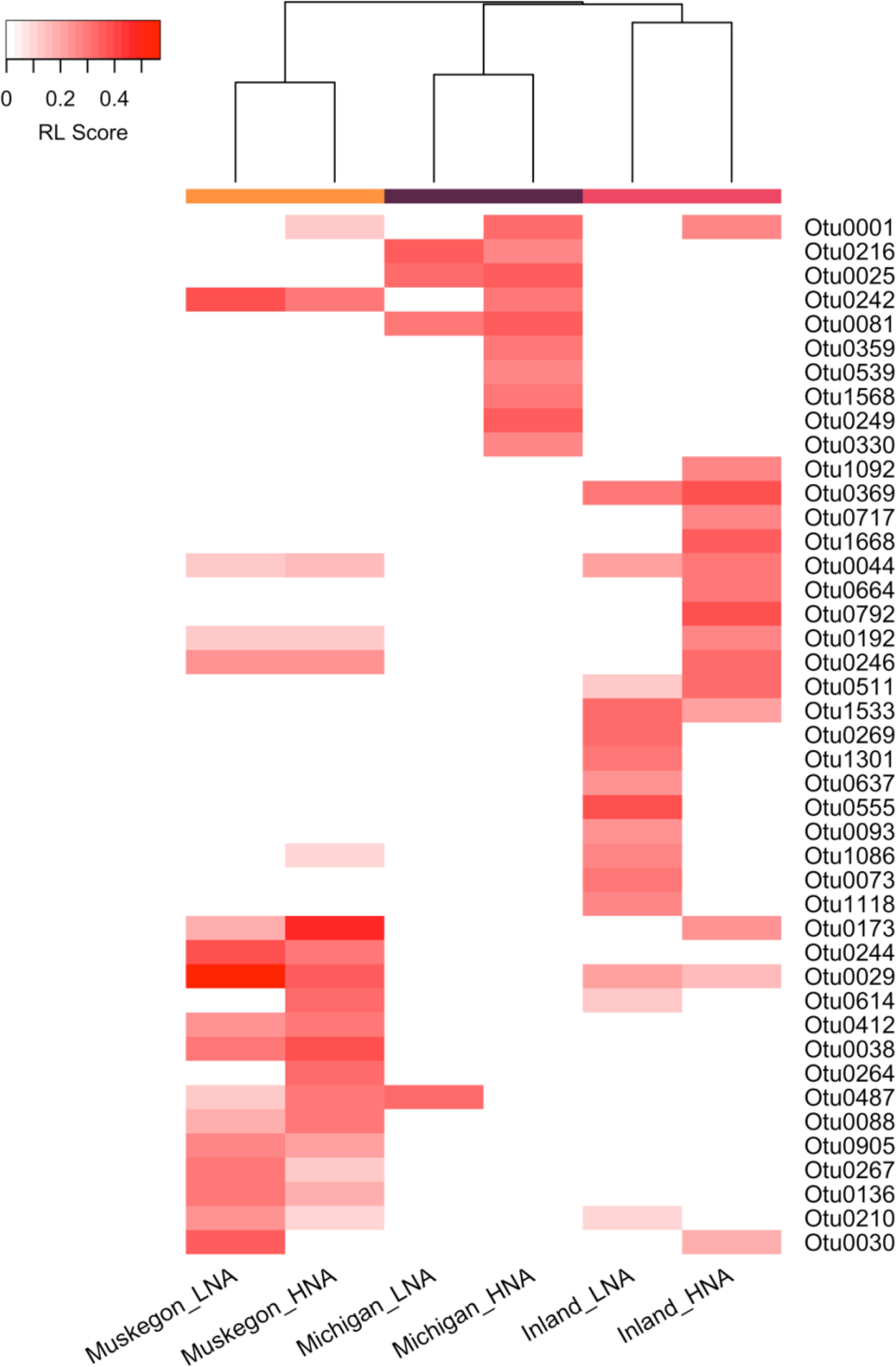
Hierarchical clustering of the RL score for the top 10 selected OTUs within each lake system and FCM functional groups with the selected OTU (rows) across HNA and LNA groups within the three lake systems (columns).

The Bacteroidetes, Betaproteobacteria, Alphaproteobacteria, and Verrucomicrobia contributed 54% of the 258 OTUs selected by the RL (Figure 5). Most selected OTUs belonging to these four phyla were associated with the LNA group (41-52% of selected OTUs), less than one third with the HNA group (14-30% of selected OTUs), and the remainder were selected as associated with both the LNA and HNA groups (23-36% of selected OTUs). In Muskegon Lake, OTU173 (Bacteroidetes;Flavobacteriales;bacII-A) was selected as the major HNA-associated taxon while OTU29 (Bacteroidetes;Cytophagales;bacIII-B) had the highest RL score for LNA OTUs. In Lake Michigan, OTU25 (Bacteroidetes;Cytophagales;bacIII-A), was selected as the major HNA-associated taxon while OTU168 (Alphaproteobacteria:Rhizobiales:alfVII) was selected as a major LNA-associated taxon. For the inland lakes, OTU369

**Figure 5:**
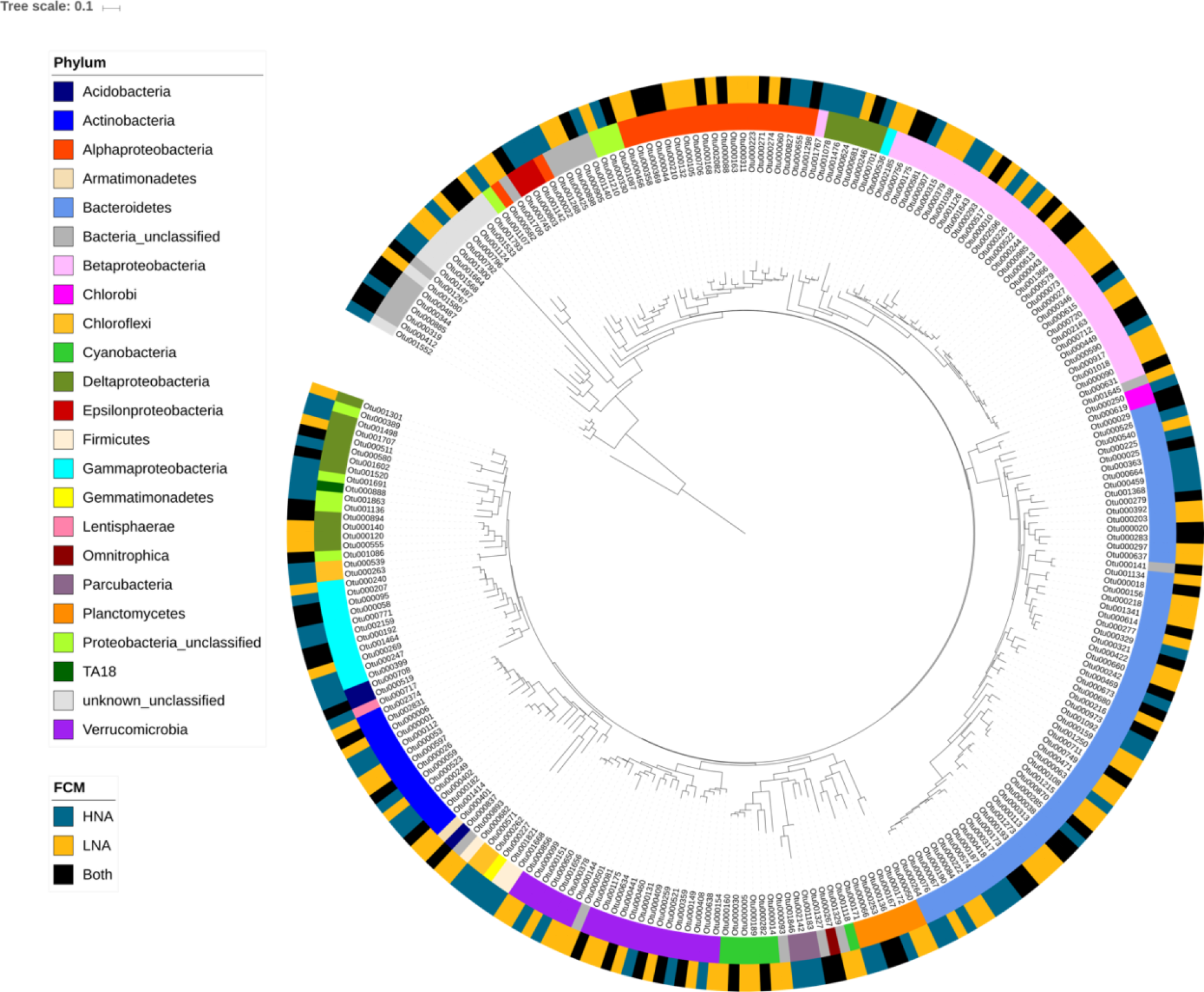
Phylogenetic tree with all HNA and LNA selected OTUs from each of the three lake systems with their phylum level taxonomic classification and association with HNA, LNA or to both groups based on the RL score threshold values.

(Alphaproteobacterial;Rhodospirillales;alfVIII) was the major HNA-associated OTU while the OTU555 (Deltaproteobacteria;Bdellovibrionaceae;OM27) was the major LNA-associated taxon. Most OTUs were selected for Muskegon Lake (153 OTUs; compared to 136 OTUs from the Inland Lakes and 20 OTUs from Lake Michigan) and 33% of these OTUs were associated with both FCM groups.

### Association with HNA and LNA is not phylogenetically conserved

To evaluate how much evolutionary history explains whether a selected taxon was associated with the HNA and/or LNA group(s), we calculated Pagel’s λ, Blomberg’s K, and Moran’s I, which are different measures for testing whether there was a phylogenetic conservation of these traits. No phylogenetic signal was detected when using Pagel’s λ with either using FCM functional group as a discrete variable (*i.e.* associating an OTU with HNA, LNA, or Both or in relation to the HNA RL score, which is a continuous variable (lambda = 0.16; P = 1) (**Figure 5**). However, there was a significant phylogenetic signal for the LNA RL score (p = 0.003, λ = 0.66), suggesting a stronger phylogenetic structure in the LNA group compared to the HNA group. This significant result in the LNA group was not found when other measures of phylogenetic signal were considered (Blomberg’s K (HNA: p = 0.63; LNA: p = 0.54), and Moran’s I (HNA: p = 0.88; LNA: p = 0.12)).

### Flow cytometry fingerprints confirm associated taxa and reveal more complex relationships between taxonomy and flow cytometric features

To confirm the association of the final selected OTUs with the HNA and LNA groups, and resolve how HNA and LNA groups correspond to OTU-level clustering of cells on the FCM fingerprints, we calculated the correlation between the density of individual small regions (*i.e.* “bins”) in the flow cytometry data with the relative abundances of the OTUs. Note that (i) as these values denote correlations, they do not indicate actual presence, and (ii) the threshold that was used to manually make the distinction between HNAcc and LNAcc (*i.e.* dotted line in Figure 6) lies very close to the border between the two regions of positive and negative correlation. OTU25 correlated with bins that when aggregated corresponded to almost the entire HNA region, whereas OTU173 was limited to bins corresponding to the bottom of the HNA region (Figure 6). In contrast, OTU369 was positively correlated to bins situated in both the LNA and HNA regions of the cytometric fingerprint, highlighting results from Figure 4 where OTU369 was selected for both HNA and LNA.

**Figure 6:**
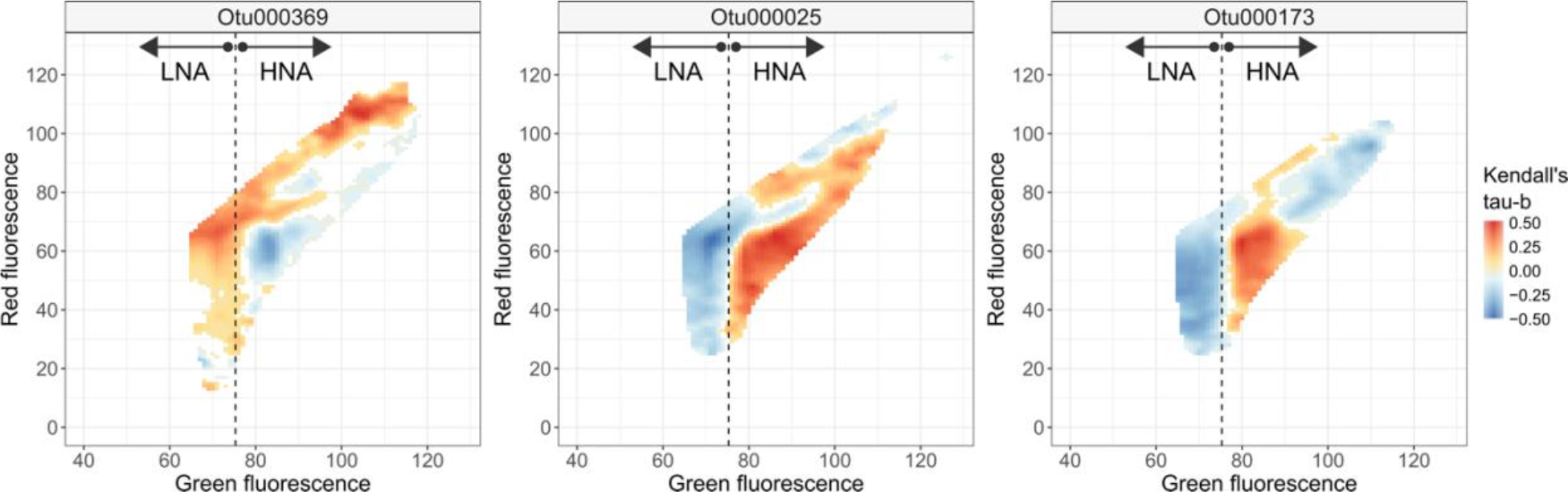
Correlation (Kendall’s tau-b) between the relative abundances of the top three OTUs selected by the RL and the densities in the cytometric fingerprint. The fluorescence threshold used to manually define HNA and LNA populations is indicated by the dotted line.

### Proteobacteria and rare taxa correlate with productivity measurements

The Kendall rank correlation coefficient was calculated between CLR-transformed abundances of individual OTUs and productivity measurements. OTU481 was the sole OTU that correlated with productivity after a multiple testing correction (Kendall’s tau-b = −0.67, P = 0.00003, P_adj = 0.016), but had a low RL score (0.022) for HNAcc and was not selected according to the Boruta algorithm. Of the top 10 OTUs selected for HNAcc according to the RL, three were still significantly associated with productivity (OTU614: P = 0.0064; OTU412, P = 0.044; OTU487, P = 0.014), but not when corrected for multiple hypothesis testing. Some OTUs that had a high RL score also had a positive response to productivity measurements, though they were insignificant after multiple testing correction. At the phylum level, only Proteobacteria were significantly correlated to productivity measurements (Kendall’s tau = 0.49, P = 0.002, P_adj = 0.05).

## Discussion

Our study furthers the integration of functional and genotypic information to determine the complex relationships between microbial diversity and ecosystem functioning. Our results confirmed previous findings that flow cytometric operational groups are distinct functional groups having divergent correlations with heterotrophic productivity. Using two machine learning based variable selection strategies, we could associate bacterial taxa identified by 16S rRNA gene sequencing to these two functional groups in three types of freshwater lake systems in the Great Lakes region. We revealed that (i) HNA and LNA cell abundances were best predicted by a small subset of OTUs that were unique to each lake type, (ii) some OTUs were included in the best model for both HNA and LNA abundance, (iii) there was no phylogenetic conservation of HNA and LNA group association and (iv) freshwater FCM fingerprints display more complex patterns related to OTUs and productivity compared to the traditional dichotomy of HNA and LNA.

Although high-nucleic acid cell counts (HNAcc) and low-nucleic acid cell counts (LNAcc) were correlated with each other, only the association between bacterial heterotrophic production (BP) and HNAcc was strong and significant. This is in line with previous reports, though past studies have focused on the proportion of HNA rather than absolute cell abundances and are strongly biased towards marine systems. For example, Bouvier et al. (11) found a correlation between the fraction of HNA cells and BP within a large dataset of 640 samples across various freshwater to marine environments (Pearson’s r = 0.49), whereas a study off the coast of the Antarctic Peninsula found a moderate correlation (R^2^ = 0.36; (17)). Another study in the Bay of Biscay also found this association (R^2^ = 0.16; (15)), however, the authors attributed this difference to be related to cell size and not due to the activity of HNA. Notably, these studies were predominantly testing the association of marine HNA groups. The high correlation coefficients observed in our study may indicate a strong coupling between freshwater carbon cycling and HNA group abundance in freshwater lake systems. Consequently, this suggests an important role of HNA bacteria in the disproportionately large role that freshwater systems in the global carbon cycle (32). It has to be noted that our study only evaluated bacterial heterotrophic production using leucine amino acid incorporation, which biases our analyses against bacterial groups that cannot import or assimilate this compound (33). Finally, as our correlations with proportional HNA group abundances also indicated less strong correlations than with absolute HNAcc, we suggest absolute HNAcc should be used to best predict and study heterotrophic bacterial production.

Similar to other microbiome studies that use machine learning, only a minority of OTUs were needed to predict the phenotype of interest, with low predictive power of each single OTU, but strong predictive capacity of the selected group of OTUs (17, 34–36). Both the RL and Boruta algorithm have been applied to microbiome studies before, for example in the selection of genera in the human microbiome associated with BMI (37), salivary pH and lysozyme activity (38), and in relation to multiple sclerosis (39) or with differing diets during primate pregnancy (40). The Boruta algorithm has also recently been proposed as one of the top-performing variable selection methods that make use of Random Forests (41). Despite the power of these approaches, improvements can be made when attempting to integrate different types of data. For example, 16S rRNA gene sequencing still faces the hurdles of DNA extraction (42) and 16S copy number bias (43). Moreover, detection limits are different for FCM (expressed in the number of cells) and 16S rRNA gene sequencing (expressed in the number of gene counts or relative abundance), therefore creating data that may be different in resolution.

The selection of different sets of HNA and LNA OTUs across the three freshwater systems indicates that different taxa underlie the universally observed HNA and LNA functional groups across aquatic systems. This is perhaps not surprising as it has been shown that there is strong species sorting in lake systems (44, 45), shaping community composition through diverging environmental conditions between the lake systems presented here (46). This high system specificity also explains the low RL scores for individual OTUs, as the spatial and temporal dynamics of an OTU diverged strongly across systems. For example, an OTU that has an RL score of 0.5 implies that on average it will only be chosen one out of two times in a Lasso model.

Some OTUs were associated with both HNAcc and LNAcc. There are multiple possible explanations for this: (a) In line with scenario 1 from Bouvier et al (11), cells transition from active growth (primarily HNA) to death or a dormant state (primarily LNA), depending on variable conditions over the spatiotemporal gradients sampled in this study. A large fraction of cells (40-95%) in aquatic systems has indeed been inferred to be dormant (47–49), in line with the predominance of LNA cells. (b) The same OTU may occur in both HNA and LNA groups due to phenotypic plasticity, which is more in line with scenario 4 from Bouvier et al (11). Bacterial phenotypic plasticity in size and morphology has been observed (50), and agrees with suggestions that HNA and LNA groups correspond to cells of differing size (12, 15, 27). (c) The association of taxa to LNA and HNA can also mean that these taxonomic groups thrive within either high or low productivity ecosystems and not necessarily that they are responsible for the change in productivity. (d) Finally, OTU level grouping of bacterial taxa can disguise genomic and corresponding phenotypic heterogeneity (51–54), which may be an alternate explanation for inconsistent associations between OTUs and FCM functional groups.

We found no clear phylogenetic conservation of association to HNAcc or LNAcc. This is in contrast to a recent study that found a clear signal at the phylum level across different aquatic systems (27). However, lake water samples were an exception to the general trend. In addition, it is notable that Proctor et al. (27) separated HNA and LNA cells based on cell size (where HNA cells were defined at approximately > 0.4 **μ**m and LNA cells were approximately 0.2-0.4 **μ**m, based on 50-90% removal of HNA cells after filtering using a 0.4 **μ**m filter), while our study separated these FCM functional groups on the basis of fluorescence intensity alone. A more direct estimation of phylogenetic conservation that directly combines cell sorting of HNA or LNA cells and sequencing, such as the approach of Vila-Costa et al. (55), will be needed to resolve these contrasting results. Considering the correlations between FCM-based phenotypic diversity and sequencing-based taxonomic diversity (56, 57), there clearly is a link between taxonomy and the structure in microbial cytometry data (17). However, the HNA/LNA dichotomy is too unresolved, as our correlation analysis between smaller regions in the cytometric fingerprint and the highly-ranked OTUs revealed a more complex relationship. This agrees with recent research, in which more than two FCM operational groups in aquatic systems were identified (17, 58, 59)7).

The Boruta algorithm and RL scores agreed on a small subset of OTUs, including the top-ranked HNA OTU for all lake systems according to RL, which motivates further investigation of the ecology of these OTUs. While little detailed information on the identities and ecology of HNA and LNA freshwater lake bacterial taxa exists, several studies identified Bacteroidetes among the most prominent HNA taxa, which is in line with our findings. Independent research by Vila-Costa et al. (55) found that the HNA group was dominated by Bacteroidetes in summer samples from the Mediterranean Sea, Read et al. (19) showed that HNA abundances correlated with Bacteroidetes, and Schattenhofer et al. (60) reported that the Bacteroidetes accounted for the majority of HNA cells in the North Atlantic Ocean. In Muskegon Lake, OTU173 was the dominant HNA taxon and is a member of the Order *Flavobacteriales* (bacII-A). The bacII group is a very abundant freshwater bacterial group and has been associated with senescence and decline of an intense algal bloom (61), suggesting their potential for bacterial production. BacII-A has also made up ∼10% of the total microbial community during cyanobacterial blooms, reaching its maximum density immediately following the bloom (62). In Lake Michigan, OTU25, a member of the Bacteroidetes Order *Cytophagales* known as bacIII-A, was the top HNA OTU. However, much less is known about this specific group of Bacteroidetes. Though, the bacII-A/bacIII-A group has been strongly associated with more heterotrophically productive headwater sites (compared to higher order streams) from the River Thames, showing a negative correlation in rivers with dendritic distance from the headwaters, indicating that these taxa may contribute more to productivity (19). In the inland lakes, OTU369 was the major HNA taxon and is associated with the Alphaproteobacteria Order Rhodospirillales (alfVIII), which to our knowledge is a group with very little information available in the literature. In contrast to our findings of Bacteroidetes and Alphaproteobacterial HNA selected OTUs, Tada & Suzuki (63) found that the major HNA taxon from an oceanic algal culture was from the Betaproteobacteria whereas LNA OTUs were within the Actinobacteria phylum.

## Conclusions

We integrated flow cytometry (FCM) and 16S rRNA gene amplicon sequencing data to associate bacterial taxa with productivity in freshwater lake systems. Our results on a diverse set of freshwater lake systems indicate that the taxa associated with HNA and LNA functional groups are lake-specific, and that association with these functional groups is not phylogenetically conserved. With this study, we show the potential and limitations of integrating flow cytometry-derived *in situ* functional information with sequencing data using machine learning approaches. This integration of data enhances our insights into which taxa may contribute to ecosystem functioning in aquatic bacterial communities. While these data-driven hypotheses will need further verification, the method is promising considering the wide application of FCM in aquatic environments, its recent application in other sample matrices (*e.g.*, faeces (64), soils (65), and wastewater sludge (66)), and the introduction of novel stains to delineate operational groups based on phenotypic traits (67).

## Materials and Methods

### Data collection and DNA extraction, sequencing and processing

In this study, we used a total of 173 samples collected from three types of lake systems described previously (46), including: (a) 49 samples from Lake Michigan (2013 & 2015), (b) 62 samples from Muskegon Lake (2013-2015; one of Lake Michigan’s estuaries), and (c) 62 samples from twelve inland lakes in Southeastern Michigan (2014–2015). For more details on sampling, please see Figure 1 and the *Field Sampling, DNA extraction, and DNA sequencing and processing* sections within Chiang et al. (46). In all cases, water for microbial biomass samples were collected and poured through a 210 μm and 20 μm bleach sterilized nitex mesh and sequential inline filtration was performed using 47 mm polycarbonate in-line filter holders (Pall Corporation, Ann Arbor, MI, USA) and an E/S portable peristaltic pump with an easy-load L/S pump head (Masterflex®, Cole Parmer Instrument Company, Vernon Hills, IL, USA) to filter first through a 3 μm isopore polycarbonate (TSTP, 47 mm diameter, Millipore, Billerica, MA, USA) and second through a 0.22 μm Express Plus polyethersulfone membrane filters (47 mm diameter, Millipore, MA, USA). The current study only utilized the 3 - 0.22 μm fraction for analyses.

DNA extractions and sequencing were performed as described in Chiang et al. (46). Fastq files were submitted to NCBI sequence read archive under BioProject accession number PRJNA414423 (inland lakes), PRJNA412983 (Lake Michigan), and PRJNA412984 (Muskegon Lake). We analyzed the sequence data using MOTHUR V.1.38.0 (seed = 777; (Schloss et al. 2009) based on the MiSeq standard operating procedure and put together at the following link: https://github.com/rprops/Mothur_oligo_batch. A combination of the Silva Database (release 123; (68)) and the freshwater TaxAss 16S rRNA database and pipeline (69) was used for classification of operational taxonomic units (OTUs).

For the taxonomic analysis, each of the three lake datasets were analyzed separately and treated with an OTU abundance threshold cutoff of at least 5 sequences in 10% of the samples in the dataset (similar strategy to (70)). For comparison of taxonomic abundances across samples, each of the three datasets were then rarefied to an even sequencing depth, which was 4,491 sequences for Muskegon Lake samples, 5,724 sequences for the Lake Michigan samples, and 9,037 sequences for the inland lake samples. Next, the relative abundance at the OTU level was calculated using the *transform_sample_counts()* function in the phyloseq R package (71) by taking the count value and dividing it by the sequencing depth of the sample. For all other taxonomic levels, the taxonomy was merged at certain taxonomic ranks using the *tax_glom()* function in phyloseq (71) and the relative abundance was re-calculated.

### Heterotrophic bacterial production measurements

Muskegon Lake samples from 2014 and 2015 were processed for heterotrophic bacterial production using the [^3^H] leucine incorporation into bacterial protein in the dark method (72, 73). At the end of the incubation with [^3^H]-leucine, cold trichloroacetic acid-extracted samples were filtered onto 0.2 µm filters that represented the leucine incorporation by the bacterial community. Measured leucine incorporation during the incubation was converted to bacterial carbon production rate using a standard theoretical conversion factor of 2.3 kg C per mole of leucine (73).

### Flow cytometry, measuring HNA and LNA

In the field, a total of 1 mL of 20 μm filtered lake water were fixed with 5 μL of glutaraldehyde (20% vol/vol stock), incubated for 10 minutes on the bench (covered with aluminum foil to protect from light degradation), and then flash frozen in liquid nitrogen to later be stored in −80°C freezer until later processing with a flow cytometer. Flow cytometry procedures followed the protocol laid out in Props et al. (56), which also uses the samples presented in the current study (Michigan and Muskegon samples). Samples were stained with SYBR Green I and measured in triplicate. The lowest number of cells collected after denoising was 2342. HNA and LNA groups were selected using the fixed gates introduced in Prest et al. (74) and plotted in **Figure S10**. Cell counts were determined per HNA and LNA group and averaged over the three replicates (giving rise to HNAcc and LNAcc). All cytometry data is available on the FlowRepository database (75): inland lakes (ID:FR-FCM-ZY9J), Michigan and Muskegon (ID:FR-FCM-ZYZN).

## Data analysis

Processed data and analysis code for the following analyses can be found on the GitHub page for this project at https://deneflab.github.io/HNA_LNA_productivity/.

### HNA-LNA and HNA-Productivity Statistics and Regressions

We tested the difference in absolute number of cells within HNA and LNA functional groups across running analysis of variance with a post-hoc Tukey HSD test (*aov()* and *TukeyHSD(); stats* R package; (76). In addition, we tested the association of HNA and LNA to each other and with productivity by running ordinary least squares regression with the *lm()* (*stats* R package; (76)).

### Ranking correlation

Ranking correlation between variables was calculated using the Kendall rank correlation coefficient, using the *kendalltau()* function in Scipy (v1.0.0) or *cor()* in R (v3.2). The ‘tau-b’ implementation was used, which is able to deal with ties. Values range from −1 (strong disagreement) to 1 (strong agreement). The same statistic was used to assess the similarity between rankings of variable selection methods.

### Centered-log ratio transform

First, following guidelines from Paliy & Shanker (77), Gloor et al. (3) and Quinn et al. (78), relative abundances of OTUs were transformed using a centered log-ratio (CLR) transformation before variable selection was applied. This means that the relative abundance *x_i_* of a taxa was transformed according to the geometric mean of that sample, in which there are *p* taxa present:

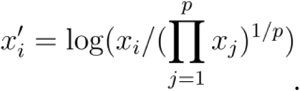

Zero values were replaced by *δ* = 1/*p*^2^. This was done using the scikit-bio package (www.scikit-bio.org, v0.4.1).

### Lasso & stability selection

Scores were assigned to taxa based on an extension of the Lasso estimator, which is called *stability selection* (28). In the case of *n* samples, the Lasso estimator fits the following regression model:

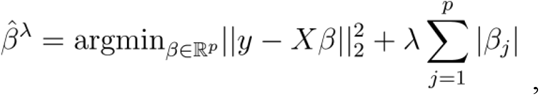

in which *X* denotes the abundance table, *y* the target to predict, which either is HNA cell abundances (HNAcc) or LNA cell abundances (LNAcc), *β* the weight of each variable and λ is a regularization parameter which controls the complexity of the model and prevents overfitting. The Lasso performs an intrinsic form of variable selection, as the weights of certain variables will be put to zero.

Stability selection, when applied to the Lasso, is in essence an extension of the Lasso regression. It implements two types of randomizations to assign a score to the variables, and is therefore also called the *Randomized Lasso* (RL). The resulting RL score can be seen as the probability that a certain variable will be included in a Lasso regression model (*i.e.,* its weight will be non-zero when fitted). When performing stability selection, the Lasso is fitted to *B* different subsamples of the data of fraction *n*/2, denoted as *X*′ and corresponding *y*′. A second randomization is added by introducing a weakness parameter α. In each model, the penalty λ changes to a randomly chosen value in the set [λ, λ/α], which means that a higher penalty will be assigned to a random subset of the total amount of variables. The Randomized Lasso therefore becomes:

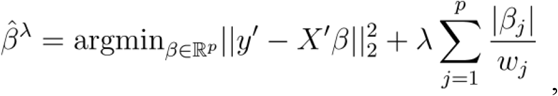

where *w_j_* is a random variable which is either α or 1. Next, the Randomized Lasso score (RL score) is determined by counting the number of times the weight of a variable was non-zero for each of the *B* models and divided by *B*. Meinshausen and Bühlmann (28) show that, under stringent conditions, the number of falsely selected variables is controlled for the Randomized Lasso when the RL score is higher than 0.5. If λ is varied, one can determine the stability path, which is the relationship between π and λ for every variable. For our implementation, *B* = 500, α = 0.5 and the highest score was selected in the stability path for which λ ranged from 10^−3^ until 10^3^, logarithmically divided in 100 intervals. The *RandomizedLasso()* function from the scikit-learn machine learning library was used ((79), v0.19.1).

### Random Forests & Boruta

The Boruta algorithm is a *wrapper* algorithm that makes use of Random Forests as a base classification or regression method in order to select all relevant variables in function of a response variable (29). Similar to stability selection, the method uses an additional form of randomness in order to perform variable selection. Random Forests are fitted to the data multiple times. To remove the correlation to the response variable, each variable gets per iteration a so-called *shadow variable*, which is a permuted copy of the original variable. Next, the Random Forest algorithm is run with the extended set of variables, after which variable importances are calculated for both original and shadow variables. The shadow variable that has the highest importance score is used as reference, and every variable with significantly lower importance, as determined by a Bonferroni corrected t-test, is removed. Likewise, variables containing an importance score that is significantly higher are included in the final list of selected variables. This procedure can be repeated until all original variables are either discarded or included in the final set; variables that remain get the label ‘tentative’ (i.e., after all repetitions it is still not possible to either select or discard a certain variable). We used the boruta_py package to implement the Boruta algorithm (https://github.com/scikit-learn-contrib/boruta_py). Random Forests were implemented using *RandomForestRegressor()* function from scikit-learn (79), v0.19.1. Random Forests were run with 200 trees, the number of variables considered at every split of a decision tree was *p*/3 and the minimal number of samples per leaf was set to five. The latter were based on default values for Random Forests in a regression setting (80). The Boruta algorithm was run for 300 iterations, variables were selected or discarded at *P* < .05 after performing Bonferroni correction.

### Recursive variable elimination

Scores of the Randomized Lasso were evaluated using a recursive variable elimination strategy (81). Variables were ranked according to the RL score. Next, the lowest-ranked variables were eliminated from the dataset, after which the Lasso was applied to predict HNAcc and LNAcc respectively. This process was repeated until only the highest-scored taxa remained. In this way, performance of the Randomized Lasso was assessed from a minimal-optimal evaluation perspective (82). In other words, the lowest amount of variables that resulted in the highest predictive performance was determined.

### Performance evaluation

In order to account for the spatiotemporal structure of the data, a blocked cross-validation scheme was implemented (83). Samples were grouped according the site and year that they were collected. This results in 5, 10 and 16 distinctive groups for the Michigan, Muskegon and Inland lake systems respectively. Predictive models were optimized in function of the *R*^2^ between predicted and true values of held-out groups using a leave-one-group-out cross-validation scheme with the *LeaveOneGroupOut()* function. This results in a cross-validated 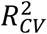 value. For the Lasso, λ was determined using the lassoCV() function, with setting eps = 10^−4^ and n_alphas = 400. The Random Forest object was optimized using a grid search where max_features was chosen in the interval 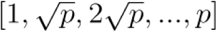 (all variables) or [1, …, *p*](Boruta selected variables) and min_samples_leaf in the interval [1, …, 5], using the *GridSearchCV()* function. The number of decision trees (n_trees) was set to 200. All functions are part of scikit-learn ((79); v0.19.1)

### Stability of the Randomized Lasso

Similarity of RL scores between lake systems and functional groups was quantified using the Pearson correlation. This was done using the *pearsonr()* function in Scipy (v1.0.0).

### Patterns of HNA and LNA OTUs across ecosystems and phylogeny

To visualize patterns of selected HNA and LNA OTUs across the three ecosystems, a heatmap was created with the RL scores of each OTU from the Randomized Lasso regression that were higher than specified threshold values. The heatmap was created with the *heatmap.2()* function (*gplots* R package) using the euclidean distances of the RL scores and a complete linkage hierarchical clustering algorithm (Figure 4).

### Correlations between taxa and productivity measurements

The Kendall ranking correlation coefficient or Kendall’s tau-b between productivity measurements and individual abundances were calculated on the phylum and OTU level using the *kendalltau()* function from Scipy (v1.0.0). P-values were corrected using Benjamini-Hochberg correction, reported as P_adj. This was done using the *multitest()* function from the Python module Statsmodels ((84); v0.5.0).

### Phylogenetic tree construction and signal calculation

We calculated the best performing maximum likelihood tree using the GTR-CAT model (-gtr - fastest) model of nucleotide substitution with FastTree (version 2.1.9 No SSE3; (85)) and visualized using the interactive tree of life (iTOL) (86). Phylogenetic signal is a measure of the dependence among a species’ trait values on their phylogenetic history (87). If the phylogenetic signal is very strong, taxa belonging to similar phylogenetic groups (*e.g.* a Phylum) will share the same trait (*i.e.* association with HNAcc or LNAcc). Alternatively, if the phylogenetic signal is weak, taxa within a similar phylogenetic group will have different traits. The phylogenetic signal was measured with both discrete (*i.e.* HNA, LNA, or both) and continuous traits (*i.e.* the RL score) using the newick tree from FastTree. For the most part, Pagel’s lambda was used (88) to test for phylogenetic signal and was calculated with the fitDiscrete() function from the geiger R package (discrete trait; (89)) and the phylosig() function from the phytools R package (continuous trait; (90)). The lambda value varies between 0 and 1, with 1 indicating complete phylogenetic patterning and 0 representing no phylogenetic patterning, leading to a tree collapsing into a single polytomy. was then used to model phylogenetic signal using Pagel’s lambda, Blomberg’s K (phylosig() function from the phytools R package (90)), and Moran’s I (abouheif.moran() function from the adephylo R package (91)).

## Supporting information

SI_Prubbens_Schmidt_etal

## Acknowledgements

PR was supported by Ghent University (BOFSTA2015000501) and MLS was supported by the National Science Foundation Graduate Research Fellowship Program (Grant No. DGE 1256260). Part of the computational resources (Stevin Supercomputer Infrastructure) and services used in this work were provided by the VSC (Flemish Supercomputer Center), funded by Ghent University, the Hercules Foundation and the Flemish Government department EWI. Flow cytometry analysis was supported through a Geconcerteerde Onderzoeksactie (GOA) from Ghent University (BOF15/GOA/006).

## Author Contributions

MLS and PR co-wrote the paper with contributions from RP, BB, NB, WW, and VJD. MLS, RP, and BB generated the data. MLS, PR, and RP performed the data analysis. PR, MLS, RP, WW, and VJD designed the study.

