## Supplementary material for "Randomized lasso associates freshwater lake-system specific bacterial taxa with heterotrophic production through flow cytometry": SI_Prubbens_Schmidt_etal

* These authors contributed equally.

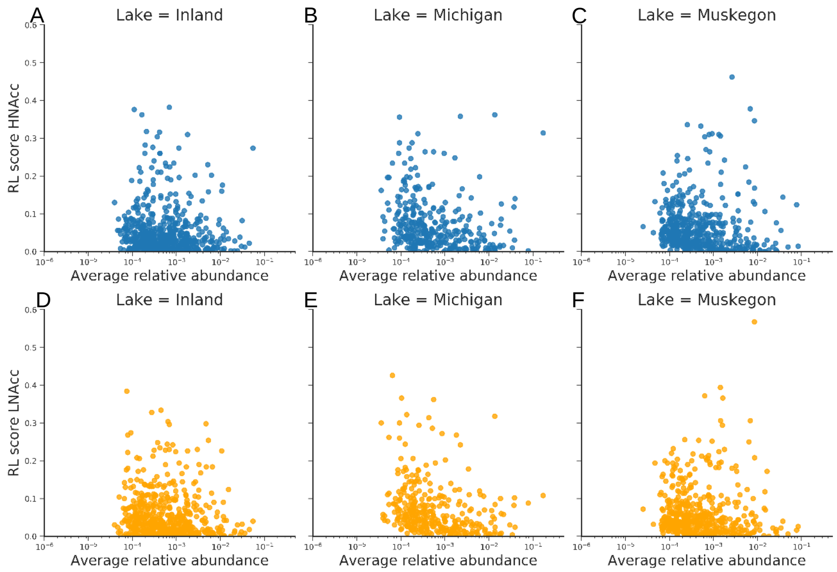


**Fig. 1.** Scatter plot of RL score versus the average relative abundance of every OTU for HNAcc (blue points, **A**, **B** and **C**) and LNAcc (orange points, **D**, **E** and **F**) for each lake system: Inland (**A** and **D**), Michigan (**B** and **E**) and Muskegon (**C** and **F**).

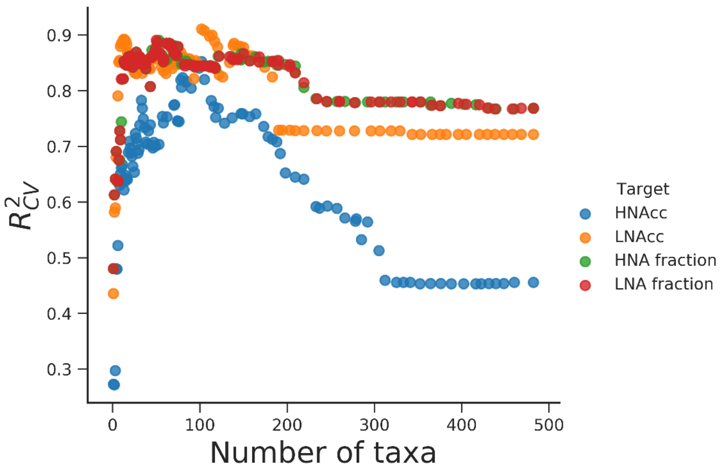


**Fig. 2.** Comparison of predictions of HNAcc and LNAcc versus relative fractions. This was done for lake Muskegon at the OTU level, expressed in terms of $R^2_{CV}$. The subset of taxonomic variables was iteratively reduced using a recursive variable elimination strategy, based on the RL score. Lowest-scored variables were removed at every step, after which the base model (i.e., the Lasso) was used to model and predict cell counts or fractions. Predictions for HNA and LNA fractions overlap (red and green dots).

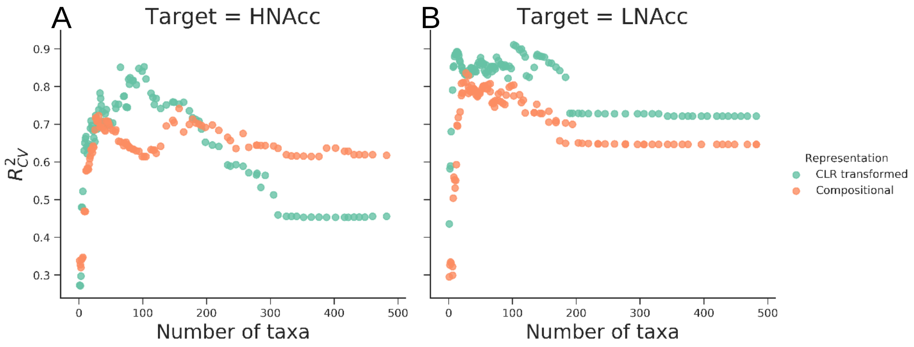


**Fig. 3.** Prediction of HNAcc (**A**) and LNAcc (**B**) for lake Muskegon at the OTU level, expressed in terms of $R^2_{CV}$ using relative abundances (compositional) and CLR transformed (CLR transformed). The subset of taxonomic variables was iteratively reduced using a recursive variable elimination strategy, based on the RL score. Lowest-scored variables were removed at every step, after which the base model (i.e., the Lasso) was used to model and predict HNAcc and LNAcc.

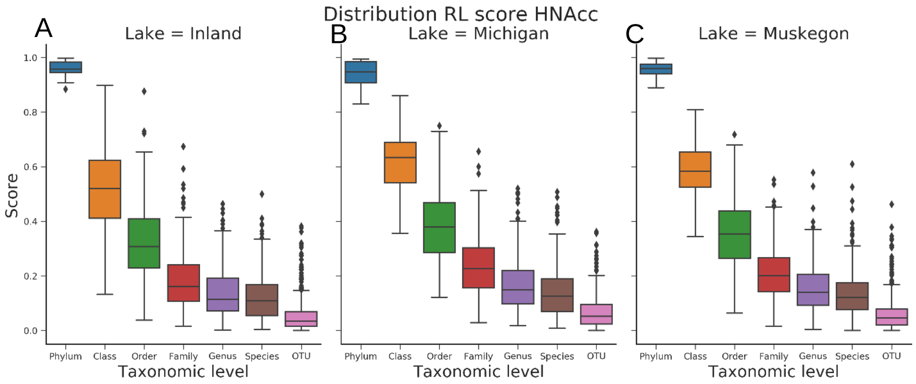


**Fig. 4.** Distribution of the RL score for all lake systems (**A**: Inland, **B**: Michigan and **C**: Muskegon) and all taxonomic levels in function of HNAcc.

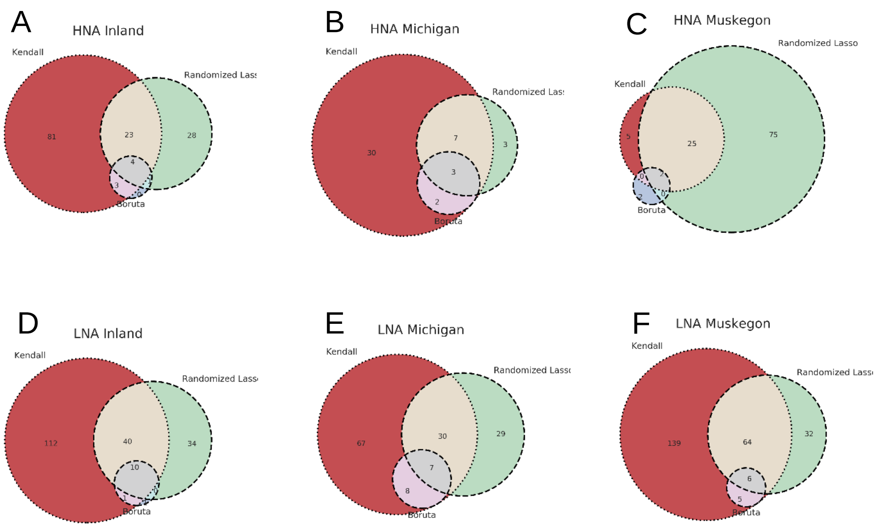


**Fig. 5.** Venn diagrams for selected OTUs according to the Kendall rank correlation coefficient, RL and Boruta algorithm. OTUs are selected for **A**: HNAcc, Inland; **B**: HNAcc, Michigan; **C**: HNAcc, Muskegon; **D**: LNAcc, Inland; **E**: LNAcc, Michigan; **F**: LNAcc, Muskegon.

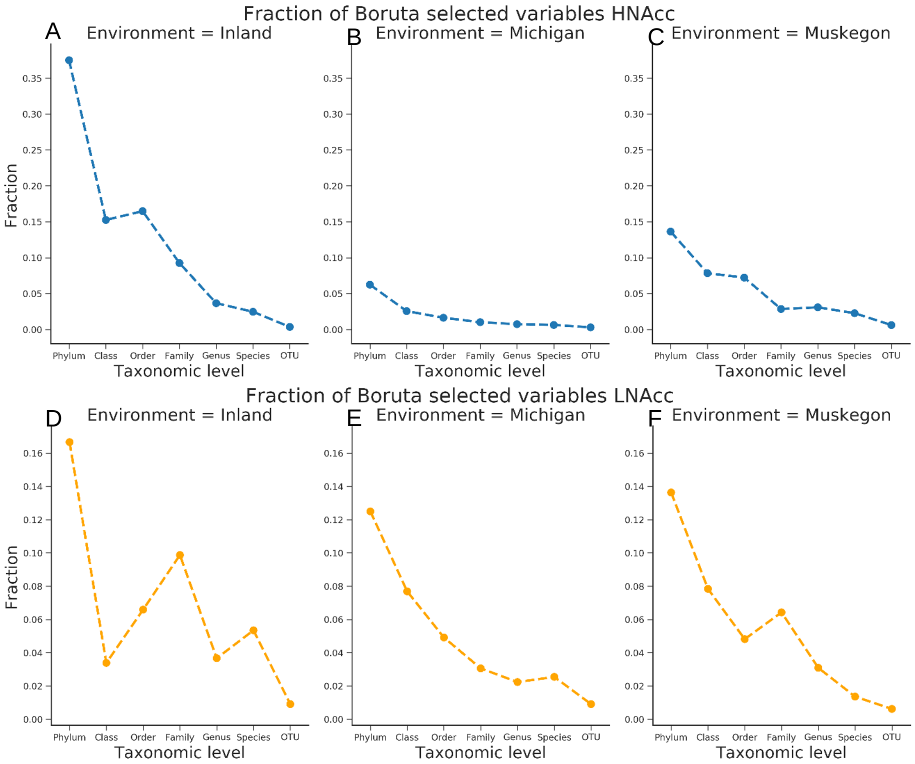


**Fig. 6.** Relative fraction of selected OTUs using the Boruta algorithm for HNAcc (blue points, **A**, **B** and **C**) and LNAcc (orange points, **D**, **E** and **F**) for each lake system: Inland (**A** and **D**), Michigan (**B** and **E**) and Muskegon (**C** and **F**).

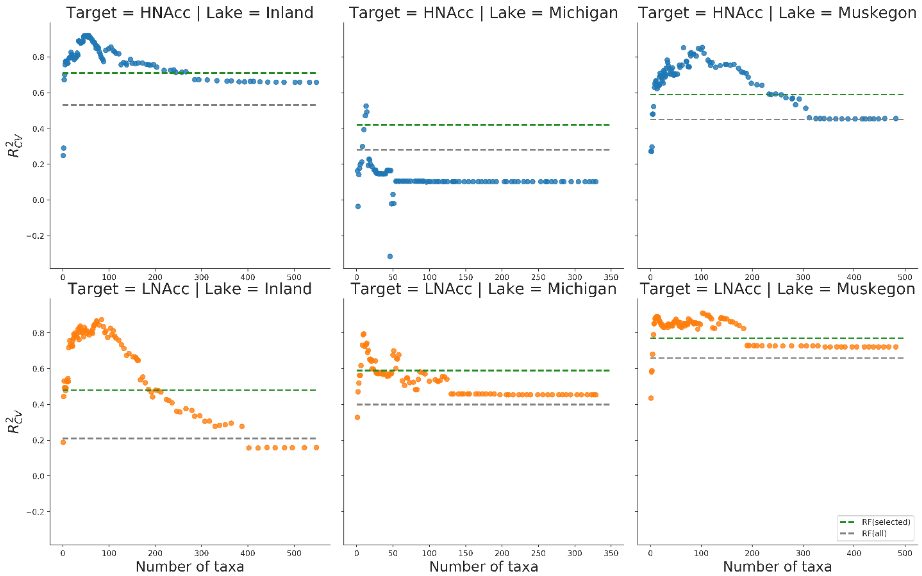


**Fig. 7.** Comparison of Random Forest predictions using all OTUs (grey dashed line) or selected OTUs (green dashed line) using the Boruta algorithm. This is compared with predictions using the Lasso and RL score at different thresholds, for HNAcc (blue points, **A**, **B** and **C**) and LNAcc (orange points, D, E and F) for each lake system: Inland (**A** and **D**), Michigan (**B** and **E**) and Muskegon (**C** and **F**). Performance is expressed in terms of $R^2_{CV}$.

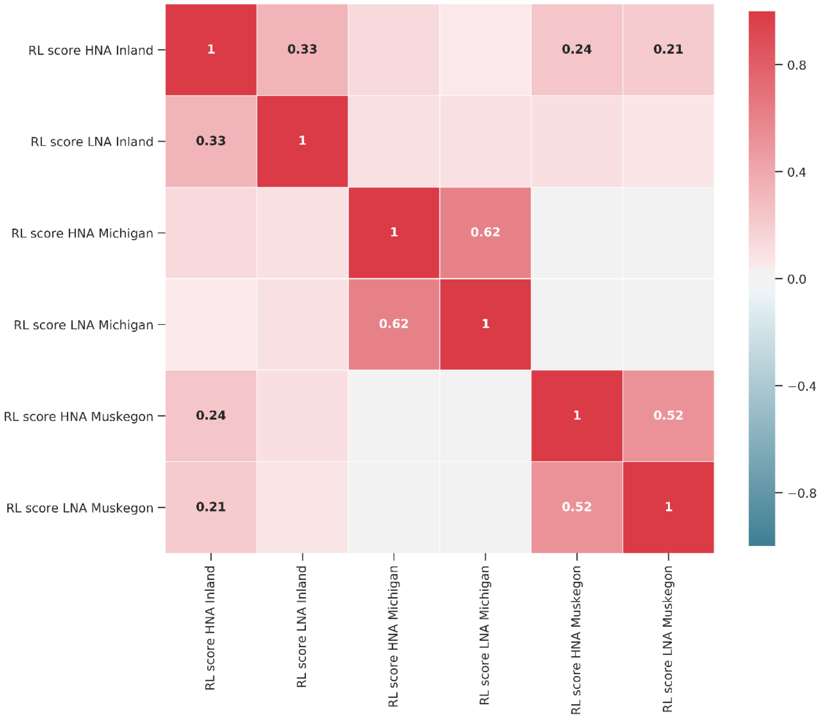


**Fig. 8.** Pearson correlations between RL scores assigned to OTUs in function of HNAcc and LNAcc between lake systems. Only those OTUs were considered that were present in all lake systems, which were 190 in total. Values are bolded if $P < 0.05$.

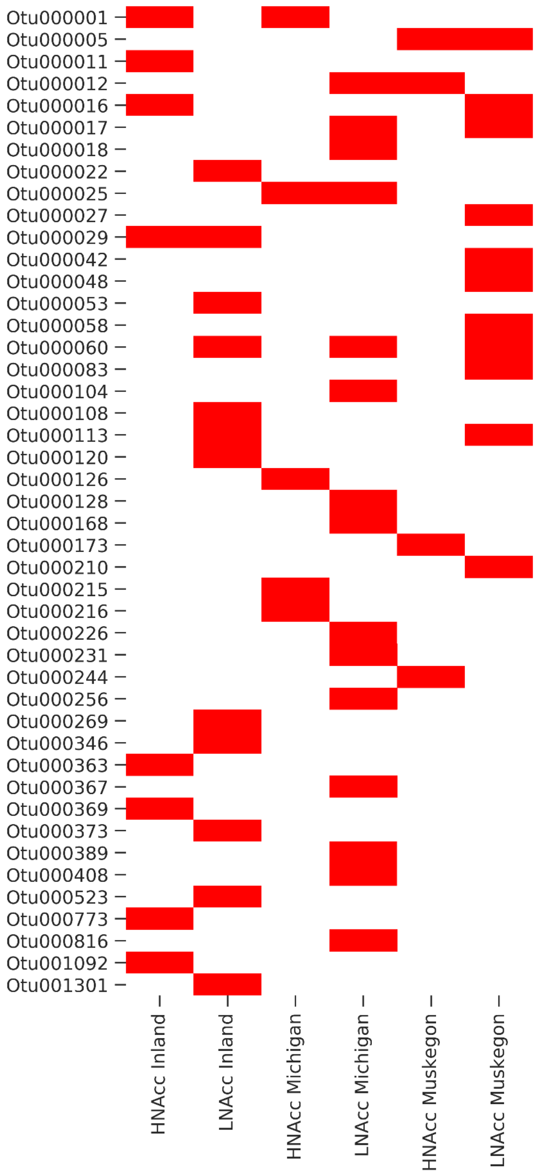


**Fig. 9.** Selected OTUs (in red) according to the Boruta algorithm for each lake system and functional group

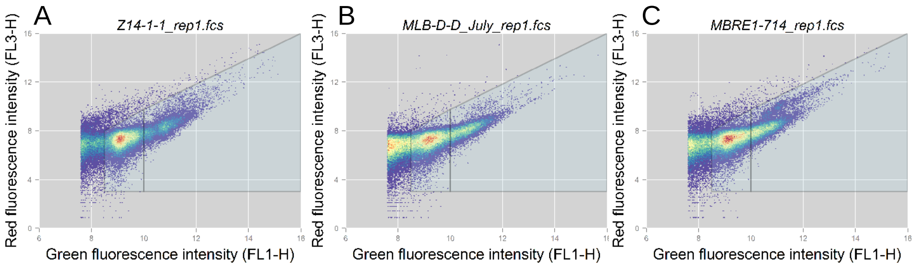


**Fig. 10.** Examples of the gating strategy to determine HNAcc and LNAcc for the three lake systems. The gating strategy is performed in the arcsinh($x$) transformed bivariate space of the FL1-H and FL3-H channel, following guidelines of Prest et al., 2013. **A**: Inland, **B**: Michigan, **C**: Muskegon.
